# Malaria parasites adjust liver stage development to synchronise the blood stage of infection with host daily rhythms

**DOI:** 10.1101/2025.11.26.690661

**Authors:** Alejandra Herbert-Mainero, Petra Schneider, Aidan J. O’Donnell, Sarah E. Reece

## Abstract

Synchronised multiplication of *Plasmodium* parasites within red blood cells causes periodic malaria fevers. Aligning blood-stage development with the vertebrate host’s feeding-fasting rhythm facilitates within-host survival and between-host transmission. We use the rodent model *Plasmodium chabaudi* to test when, following development in the liver, the blood stage of infection begins. We find egress from the liver into the blood is aligned with the time of day of rhythmic host feeding, but only in wild type hosts, with egress occurring after a fixed period of pre-erythrocytic development in hosts without a functional canonical clock. However, perturbing the duration over which parasites enter the bloodstream does not affect their multiplication rate in the first few IDCs, suggesting fitness benefits from timing egress anticipates rhythmic challenges or opportunities later in the infection.

## Introduction

The daily rotation of the Earth exposes organisms to predictable environmental cycles that impact on fitness. Endogenous time-keeping mechanisms, including circadian clocks, are thought to have evolved to enable organisms to appropriately schedule activities to align with these environmental rhythms^1,2,3^. While the endogenous clocks of free-living organisms are the subject of much research; whether, how, and why parasites keep time is poorly understood^4^. Parasites (including pathogens and microbes) are exposed to the daily rhythms expressed by hosts and vectors, including rhythms in immune responses, behaviours, and nutrient availability that follow from hosts feeding during their active phase. Many parasites have complex life cycles, using multiple hosts/vectors, and must cope with physiological rhythms in their current host while also anticipating rhythms in the next host that dictate the time of day at which transmission occurs^5^. For example, malaria parasites (*Plasmodium spp.)* rhythmically invade and burst the red blood cells (RBCs) of their vertebrate host. This intraerythrocytic development cycle (IDC) lasts for multiples of 24hrs, and is timed to best exploit the nutrients parasites need from the host’s food while also ensuring their transmission forms have matured by night time when mosquito vectors bite^5–13^. Alignment is actively maintained by a parasite circadian oscillator; when host feeding rhythms are experimentally shifted, *Plasmodium chabaudi* shortens the IDC by 2-3 hours per cycle to regain alignment with host feeding rhythms^9–12,14–19^. Such plasticity comes with a cost to per-cycle productivity^9,12^, although completing additional rounds of multiplication during rescheduling can compensate for this cost^18,20^.

Like *Plasmodium*, many parasites reside in multiple tissues within each host/vector, and therefore experience a variety of host tissue-specific rhythms during their life cycle, that provide time of day-specific opportunities to exploit or dangers to evade. Before the IDC begins, malaria parasites enter the vertebrate host during the bite of an infected mosquito and subsequently undergo extensive replication in hepatocytes. Once inside a liver cell, these exoerythrocytic forms (EEFs) multiply by several orders of magnitude more than blood stages and are intensely nutrient-demanding^21–24^. Because we expect that the circadian clock used during the IDC^12,14–19^ will also be ticking in EEFs, and because hepatocytes supply nutrients rhythmically^10,11,25^, temporal alignment to host feeding might benefit EEFs. Alternatively, because replication of nuclei within EEFs occurs on a time scale much faster than 24 hours^26–30^ and EEFs can alter hepatocyte transcription to secure resources^31–33^, this may relax any circadian constraint on nutrient acquisition by EEFs. In contrast, altering host cell transcription to maximise nutrient accessibility is not possible in mammalian RBCs (which lack a nucleus), forcing IDC stages to be more reliant on exploiting host feeding-fasting rhythms than EEFs. While karyokinesis, in both blood and liver schizonts, is asynchronous, the subsequent organellar and membrane biosynthesis in the multinucleated schizont as well as segregation of progeny by cytokinesis is fairly synchronous^26–30,33–38^. This opens avenues for EEFs to either synchronise cytokinesis or egress and start the IDC in alignment with host rhythms. Whether parasites schedule egress ‘on time’ to benefit the IDC, or whether EEFs are unable to synchronise cytokinesis, or delay egress, forcing IDC stages to synchronise with each other and host rhythms once in blood, are open questions.

Here, we ask whether *P. chabaudi* has evolved to start the blood stage of infection synchronously and aligned to host feeding-fasting rhythms by experimentally testing the following hypotheses. **Hypothesis 1**: parasites egress using time of day cues directly derived from host feeding-fasting rhythms as observed for IDC stages. For example, responding to isoleucine rhythms would allow parasites to maximise fitness by ensuring the IDC is appropriately aligned to host feeding-fasting rhythms from the point of initiating the blood stage^25^; **Hypothesis 2**: egress cues are derived from host TTFL-driven rhythms (the transcription-translation-feedback-loop that runs the canonical circadian clock, which is most responsive to light), and so indirectly reflecting the timing of host feeding-fasting rhythms.

While this allows parasites to align egress with host feeding in general, natural perturbations to feeding schedules would cause misalignment, resulting in fitness costs. With considerable variation in the time-of-day infectious mosquito bites are received^39–42^, causing for example dawn and dusk bites introducing parasites 12 hours apart, both hypothesis 1 and 2 require the duration of exoerythrocytic development to be plastic. **Hypothesis 3**: parasites may have evolved a fixed duration of exoerythrocytic development, which matches the duration between the average time of day that infectious bites are received and the average time of day that hosts feed. This strategy ensures that egress usually coincides with host feeding-fasting rhythms, and reduces the costs of detecting and interpreting time-of-day cues, but natural variation in mosquito biting time of day^39–42^ or the host’s feeding schedule would cause misalignment of IDC stages and fitness costs. **Hypothesis 4**: EEFs may be entirely unresponsive to host rhythms, completing development in an unsynchronised manner, and egress over a prolonged duration. This may be caused by differences in the rates of development or productivity (number of progeny produced) between individual EEFs within a host, and could even be advantageous if, for example, gradual release into the blood avoids stimulating dangerous immune responses. We test these four hypotheses by manipulating when *P. chabaudi* infections are initiated by mosquito bite with respect to the phase of host feeding-fasting and TTFL-driven rhythms, using wild type and TTFL-deficient mice, and we also investigate the fitness consequences of initiating the blood stage in different manners.

## Results

Our *in vivo* model system allows us to manipulate the host’s feeding-fasting rhythms and/or TTFL-driven rhythms (i.e. the presence of a functional canonical circadian clock), relative to the time of day at which *P. chabaudi* (genotype AJ) infections are initiated, while also allowing parasites to respond to the full complexity of the daily rhythms and physiological processes they encounter inside a mammalian host. We carried out three experiments to determine if the blood stage of infection is initiated in a rhythmic manner, when this occurs and in response to which host rhythms, and whether starting the IDC synchronously maximises blood stage multiplication rates.

### The influence of host rhythms on parasite egress from the liver

We asked whether patterns of accumulation in the blood are determined by either host feeding-fasting (hypothesis 1) or TTFL-driven (hypothesis 2) rhythms, or are unresponsive to host rhythms, which can occur either because the duration of exoerythrocytic development is fixed (hypothesis 3), or because egress occurs over a prolonged window (hypothesis 4). To distinguish between these four hypotheses, we initiated infections by mosquito bites in hosts whose TTFL-driven rhythms, feeding-fasting rhythms, or both were disrupted and monitored the accumulation of blood stage parasites as a proxy for egress (Alignment of egress experiment, Figure 1a). We manipulated the presence of TTFL-driven host rhythms by using either wildtype C57BL/6 mice in a standard 12:12hr photoschedule (WT) or *Per1/2* null TTFL-deficient mice in constant darkness (KO) that are arrhythmic when kept under constant conditions (constant darkness and *ad libitum* (AL) feeding)^43^. Experimentally controlled feeding schedules (restricted feeding, RF) allowed us to vary the timing of host feeding-fasting rhythms relative to the time of day of infection in wildtype mice, and to create a feeding-fasting rhythm in the absence of TTFL-driven rhythms by using the *Per1/2* null mice.

**Figure 1.**
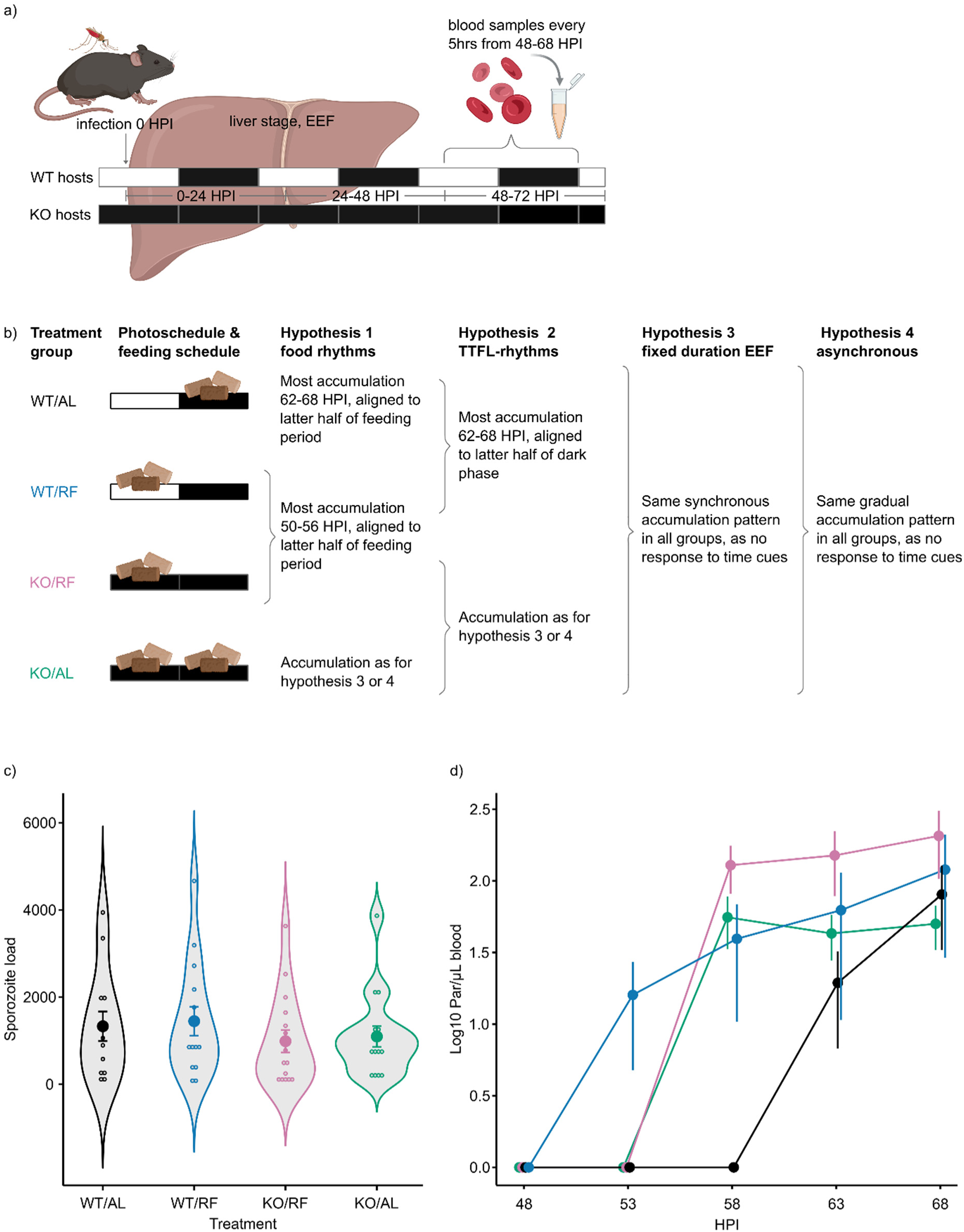
Alignment of egress experiment. **a)** Experimental setup. *Plasmodium chabaudi* AJ liver stage infections were initiated in wildtype mice in a standard 12hr light (white):12hr dark (black) photoschedule (WT) or in *Per1/2* null mice in continuous darkness (KO) as denoted by the horizontal bars for each host type. **b)** Treatment groups and predicted outcomes for the 4 hypotheses. Mice experienced different feeding-fasting rhythms, with food pellets representing feeding windows. WT hosts fed *ad libitum* (AL) eat in the dark phase but KO mice in continuous darkness and fed *ad libitum* (AL) eat continuously. Restricted feeding for 12 hours (RF) decouples the alignment between feeding-fasting and TTFL-driven rhythms in WT mice, and creates a feeding-fasting rhythm in KO mice^12,44^. The grouping of predicted patterns of parasite accumulation in the blood is shown for each of the 4 hypotheses. **c)** Violin plot for sporozoite loads, i.e. the sum of residual sporozoites in the salivary glands of all mosquitoes that fed on a host. Datapoints show sporozoite loads per host, with mean ± sem shown for each treatment group. **d)** Log_10_ mean *Plasmodium chabaudi* AJ densities per µL blood ± sem at 5-hourly intervals between 48-68 HPI for n=8 infected (out of n=15 exposed) WT/AL mice; 9 (15) WT/RF; 9 (16) KO/RF and 6 (16) KO/AL mice. HPI: hours post infection by mosquito bite; colours represent the different treatment groups.

We predicted that patterns of parasite accumulation in the blood across the 4 treatment groups should vary between hypothesis 1-4 (Figure 1b) as follows. Under hypothesis 1, we expect synchronous accumulation during the latter part of the feeding window for all groups with a feeding-fasting rhythm (WT/AL, WT/RF, KO/RF). Specifically, egress should occur towards the end of the dark/feeding period for the WT/AL group, with this pattern shifting by 12 hours in both RF groups to match the hosts’ feeding windows. Under hypothesis 2, both WT groups should follow identical, and synchronous, patterns of accumulation towards the end of the dark period, independent of host feeding-fasting schedules, whereas the KO groups should display the same pattern as each other, which is likely different from that of the WT groups. Under hypothesis 3, all groups should display the same and fairly synchronous pattern of accumulation due to a fixed duration of exoerythrocytic development. Whilst mosquito infections naturally occur at night, we infected hosts during the light phase (according to the photoschedule of WT hosts) to ensure the expected time of egress under hypothesis 3 is decoupled from the dark/light cycle implicated by hypothesis 2. Finally, observing gradual accumulation over a prolonged window in all four groups would support hypothesis 4.

We initiated infections by exposing each mouse to the bites of 24 mosquitoes (54% *P. chabaudi* infection prevalence) at 11am GMT, which is 4 hours after lights on (Zeitgeber Time, ZT4) for the WT hosts (Figures 1a, S1). Sporozoite exposure was consistent across treatment groups (F_3,56_=0.558, p=0.645, Figure 1c). The relatively low sporozoite load of 1205 ± sem 143 sporozoites per mouse resulted in 52% ± sem 6% of exposed hosts becoming infected, with equal prevalence between treatment groups (χ^2^(3)=2.004, p=0.572). Total parasite densities achieved at 68 HPI did not differ between treatment groups (F_3,28_=2.072, p=0.127), which – assuming the majority of parasites have egressed – suggests that host rhythms do not influence EEF productivity. However, the dynamics of parasite accumulation varied between treatment groups (HPI by Treatment interaction: χ^2^(9)=29.87, p<0.001; Figure 1d) indicating variation in the timing of egress. This eliminates hypotheses 3 and 4 because they predict egress patterns to be the same across all treatment groups. Instead, the observed patterns support a response to host feeding-fasting (hypothesis 1) or TTFL-driven rhythms (hypothesis 2) (Figure 1b). Specifically, despite the feeding window being offset by 12 hours in the WT groups, parasites are first detected in the blood during the latter half of the feeding window for both these groups (WT/AL, 63HPI; WT/RF, 53 HPI), supporting hypothesis 1. However, a role of TTFL-driven rhythms (hypothesis 2) is also suggested because both KO/RF and KO/AL groups follow the same pattern of accumulation as each other (parasites first detected in the blood at 58 HPI), which differs from the WT groups.

A combination of hypotheses 1 and 2 is plausible if the host’s canonical clock is necessary for parasites to egress in alignment with host feeding-fasting rhythms. This is consistent with parasites exhibiting a fixed duration of exoerythrocytic development which results in synchronous egress between 53-58 HPI (Figure 1d) in the absence of host TTFL-driven rhythms, as observed in the KO groups. With *Anopheles* mosquitoes usually blood feeding and consequently initiating infections during the mid part of the night, egress after a fixed developmental duration of 53-58 HPI (revealed in the KO groups) would coincide with the end of the feeding period in normal circumstances (i.e. in WT/AL hosts). In this ‘Alignment of egress’ experiment we deliberately shifted the time of infection by mosquito bite to the light phase for WT hosts in order to better distinguish between our hypotheses. Therefore, to test whether egress normally occurs at the end of the dark/feeding phase, we performed a second experiment (Time of egress experiment) where we infected WT/AL hosts by mosquito bite during the dark phase (Figure S1) and observed that parasites start accumulating in the blood at the end of the feeding window/dark phase (54 HPI, Figure 2). Because this is considerably earlier than in the WT/AL group in the ‘Alignment of egress’ experiment (63 HPI, Figure 1d) it confirms that those parasites were plastically extending the duration of exoerythrocytic development to initiate the IDC in alignment with the host’s feeding-fasting rhythms.

**Figure 2.**
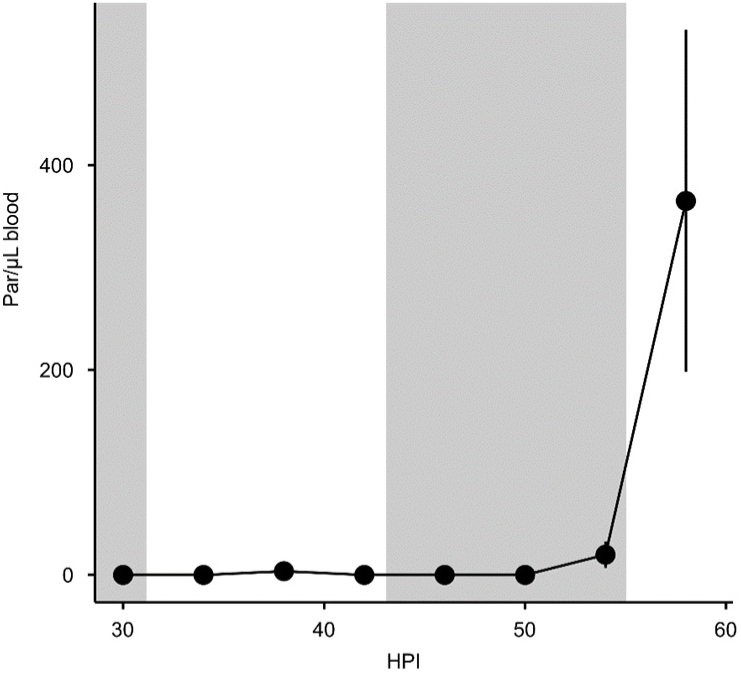
Time of egress experiment. Mean *P. chabaudi* AJ densities per µL blood ± sem at 4-hourly intervals between 30-58 HPI for n=5 infected (out of n=13 exposed) WT/AL hosts infected by mosquito bite 5 hours into the night time. The shaded areas represent darkness during the 12hr light:12hr dark photoperiod.

### Fitness consequences of starting the first IDC synchronously

Having found support for parasites in normal circumstances (i.e. WT/AL hosts) adjusting their egress schedule in response to the timing of host rhythms, we tested the fitness consequences of this strategy with a ‘Simulated egress’ experiment. We predicted that egress over a defined window towards the end of the feeding/dark phase is optimal for parasites because it allows IDC stages to garner fitness benefits from developing in alignment to host feeding-fasting rhythms (“extrinsic fitness benefits”^9,18^) and there may also be “intrinsic fitness benefits” from synchronous development. For example, temporally compartmentalising DNA replication and metabolism is an assumed intrinsic benefit of using circadian clocks to schedule these processes.

To test whether the pattern of egress affects the multiplication rate of IDC stages, we experimentally simulated “synchronous” versus “spread-out” egress patterns. Infections were either started with a single inoculum of IDC parasites at ring stage aligned to host feeding rhythms, or with three injections of equal numbers of ring stages spaced 8 hours apart in which one inoculum was aligned to host feeding rhythms (Figure 3a). Furthermore, we also tested whether the consequences of starting the IDC in a synchronous or spread-out manner depend on whether hosts have a functional (WT/AL hosts) or disrupted (KO/AL hosts) TTFL clock. We predicted that if parasites in WT hosts multiply faster when egress is synchronous compared to spread-out they garner extrinsic benefits and that if this difference occurs in KO hosts, they additionally or alternatively garner intrinsic benefits.

**Figure 3.**
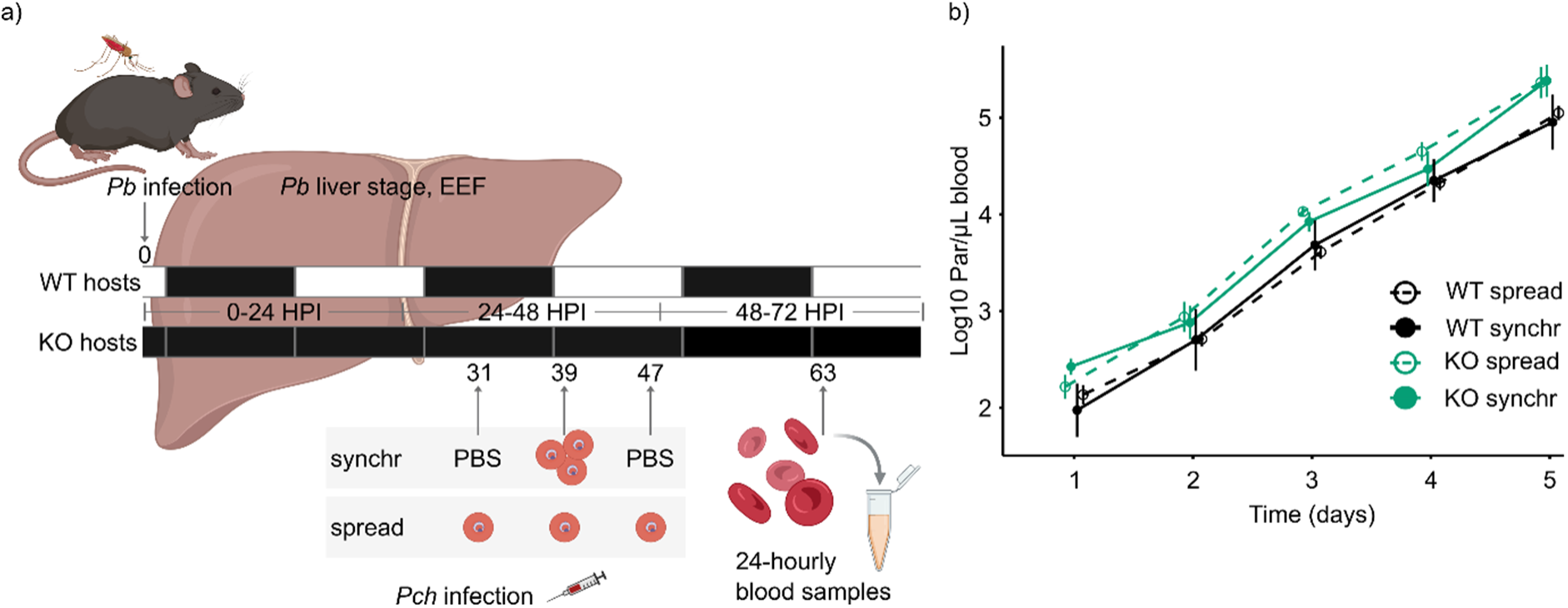
Simulated egress experiment. **a)** Experimental design testing for fitness consequences of synchronous (synchr) or spread-out (spread) establishment of the IDC. Wildtype (WT, n=10 per group) and *Per1/2* null (KO, in constant darkness, n=5 per group) mice were exposed to *P. berghei* (Pb) infected mosquitoes, followed by injection of 3*10^5^ *P. chabaudi* (Pch) ring stage infected RBCs to initiate the IDC. Inoculation regimes are “spread” (10^5^ parasites each at 31, 39 and 47 HPI) or “synchr” (3*10^5^ parasites at 39 HPI and PBS placebo at 31 and 47 HPI). **b)** Log_10_ mean *P. chabaudi* AJ densities per µL blood ± sem for spread-out (dashed) and synchronous (solid) simulated egress strategies from day 1 (63 HPI) to day 5 (159 HPI) after exposure to *P. berghei* infected mosquitoes, in WT (black) and KO (green) hosts. HPI= hours post infection by mosquito bite. GMT= Greenwich Mean Time.

We infected WT (in a 12:12h light-dark photoschedule) and KO (in constant darkness) hosts with *P. berghei* by mosquito bite to initiate a liver stage infection before receiving *P. chabaudi* blood stage parasites via injection (Figure 3a, Figure S1). We pre-infected hosts to maximise ecological relevance, such that *P. chabaudi* established its blood stage infection in hosts experiencing a liver stage infection. Quantifying *P. chabaudi* using a species-specific quantitative PCR that does not detect *P. berghei* parasites, prevented any liver-egressed parasites from confounding our results ^45^. Each mouse was exposed to on average 4.0 ± sem 0.2 *P. berghei*-infected mosquitoes, with no differences between treatment groups (F_3,26_=0.592, p=0.626). From 31 to 47 hours after *P. berghei* infections were initiated, each host received three injections, spaced 8 hours apart (Figure 3a). To simulate synchronous egress, *P. chabaudi* ring stages were administered only in the middle injection, timed to align the IDC to host rhythms for the WT hosts (a few hours into the egress window, i.e. 1h after lights on, ZT1), and the same number of ring stage parasites were split equally across all three injections to simulate spread-out egress (Figure 3a, Figure S1).

Over the 5 days following *P. chabaudi* infection, neither densities (χ^2^ (3)=2.348, p =0.504) or multiplication rates (5.8-fold growth/day) of parasites differed between treatment groups (Day by Treatment interaction: χ^2^(12)=7.454, p=0.826; Figure 3b). Thus, starting the IDC synchronously or spread over 16 hours does not impact parasites’ ability to establish a blood stage infection in rhythmic or arrhythmic hosts, suggesting that parasites do not receive extrinsic or intrinsic benefits from rhythmic egress, at least during the first IDCs.

## Discussion

We conducted two experiments to test whether *P. chabaudi* egresses from the liver and starts the IDC in alignment with the host’s feeding-fasting rhythms that also optimise the development and multiplication of blood stage parasites. We tested four hypotheses and do not find clear, unequivocal support for any of them individually. However, our results are consistent with a scenario in which parasites adjust the duration of liver stage development to align egress with host feeding-fasting rhythms, but can only achieve this when the host has a functional canonical (TTFL) circadian clock. The difference in the timing of blood stage accumulation in the WT/AL and WT/RF hosts is unlikely to be due to merely disrupting the alignment between rhythms driven by the light-dark cycle and by feeding-fasting. If internal desynchrony was the cause, parasites in KO/RF hosts would have initiated the blood stage at the end of the feeding window because feeding-fasting provides the dominant time of day information in KO hosts (in constant darkness). Our third experiment probed whether highly synchronous egress, timed to align with the host’s feeding-fasting rhythms, enhances the multiplication rate of IDC stages. Contrary to expectations, the multiplication rate during the first few IDCs was similar for synchronous or temporally dispersed (over 16 hours) initiation of the blood stage. Studies that determine the developmental duration from the liver to the blood stage of infection are sparce, especially for *P. chabaudi*. Our data indicate that *P. chabaudi* EEFs typically egress 53-58 HPI, consistent with previous estimates of 52-53 hours^46^, and we also demonstrate, for the first time, that this duration can be plastically adjusted to align egress with host feeding. Altering the time of infection by mosquito bite for unperturbed WT hosts between the ‘Alignment’ and ‘Time of egress’ experiments confirms that egress starts at the end of the host feeding phase (Figure 2). In addition, following a feeding window shift of 12 hours, parasites egressed ∼10 hours earlier, which is as close to 12 hours as our 5-hourly sampling schedule allows (Figure 1d).

The signals used by parasites to end the liver stage and begin the blood stage of infection are unknown. Parasites could use time of day cues sensed at the point of infection (or after) to set the schedule for the development of EEFs, or EEFs could complete development and wait for a time-of-day signal (a “gate”) to egress, or use a combination of both strategies.

The different accumulation patterns in the WT and KO/RF groups suggests that the cues for egressing ‘on time’ involve the canonical (TTFL) circadian host clock. This was unexpected because IDC stages use cues from only the host’s feeding-fasting rhythm (potentially, blood isoleucine concentration^25^) to set their schedule. Fundamental differences between EEFs and IDC stages in nutrient acquisition and host cell environment may explain EEF’s dual dependence on food intake and the TTFL. EEFs have no haemoglobin to digest, are highly dependent on lipid synthesis, and do not appear to have the new permeability pathways (NPPs) that facilitate the increased isoleucine uptake by infected erythrocytes^33,47,48^.

Furthermore, rhythmic cues may differ in level and amplitude between the hepatocytes and RBCs, and EEFs may require higher amplitude rhythms compared to IDC stages due to their nutrient-rich environment. Combining a TTFL- and food-driven response may result in more reliable egress timing when the host eats at an unusual time at the day, or for hosts under energetic stress that have an altered feeding-fasting rhythm. Thus, EEF egress may be governed by both TTFL and feeding cues, analogous to the dual cue regulation proposed for the conversion of *Plasmodium* IDC stages into sexual transmission stages^49,50^. More generally, responding to dual cues facilitates the perception of noise^50^ and provides a more robust response in fluctuating environments. Furthermore, the dampening of feeding-imposed liver rhythms in TTFL-deficient hosts^51^ could also explain why egress was not aligned to the timing of feeding in KO/RF hosts. For example, the rhythmicity of many lipid species, glucose, glycogen and triglycerides requires the presence of either the central TTFL clock or the peripheral liver TTFL clock^25,51–56^ and hormones like insulin and glucocorticoids act as a systemic entrainer of hepatic rhythms to feeding^57–59^. Daily oscillations in hepatocyte translation efficiency, protein accumulation, and cell size – processes that peak at the start of the fasting phase (light phase in mice) but depend on the alignment of photoschedule and feeding rhythms^60^ – may further erode liver rhythms in KO hosts, preventing parasites from time-keeping. These oscillations are absent in mature erythrocytes and so a reliance on the TTFL-driven aspects would be a constraint in IDC stages, perhaps explaining the lack of TTFL-dependency in IDC rhythmicity.

We also investigated the fitness benefits for parasites of adopting a plastic strategy for timing the initiation of the blood stage of infection. The evolution of adaptive plasticity requires a predictably fluctuating environment which can be reliably assessed, such as a repeatable environmental rhythm. Additionally, the energetic costs associated with sensing and responding to environmental cues must be outweighed by the fitness benefits of adopting a context-specific phenotype. Contrary to our expectations that a tightly synchronised egress, aligned to host feeding-fasting rhythms, would be favoured, a variable duration of the liver stage did not alter EEF productivity (Alignment of egress experiment).

Although low parasite densities may obscure subtle group differences, our multi-IDC simulated egress experiment also showed that a synchronous start conferred no advantage to parasites in WT or KO hosts. This agrees with predictions that (IDC) schizonts burst when maximum size / progeny number is reached rather than after a specific duration of development^29,30,38^. Applying this strategy to EEFs suggests that EEF rhythmicity may not gain a fitness advantage through increased productivity, but that the timing/synchrony of egress could be adaptive. Putative benefits of starting the blood stage infection at a specific time of day could include overwhelming immune defences, evading peak immune activity. or minimising nutrient limitation. These factors may be negligible during the first IDCs when parasites densities remain low, but setting the right IDC schedule early ensures parasites are ‘on time’ when immune responses and nutrient limitation develop due to rapidly increasing parasite densities. Repeating the simulated egress experiment by starting infections with a higher parasite density, in semi-immune or nutrient limited hosts is necessary to test whether the fitness benefits of egressing at the end of the host’s feeding window become apparent when infections progress. Furthermore, our approach for simulating spread-out egress may not have resulted in enough parasites being out of alignment with host rhythms to detect fitness costs. Parasite densities at the start of the infections were low and. while parasites in the first and third inoculums were misaligned by 8 hours, this may not be enough to significantly reduce replication^18,20^. Alternatively, *P. yoelii* and *P. berghei* EEFs egress spans 6-10 hours, similar to our observations in *P. chabaudi*^23,61^. Such intermediate levels of synchrony could, as predicted by theory diffuse nutritional demand over a broader interval, thus moderating competition between genetically related parasites within the host^62,63^. Our fully synchronous and spread-out treatments may therefore be equidistant from the true optimum of intermediate synchrony, obscuring fitness differences. Thus, the best strategy may be for parasites to use host time of day cues to plastically adjust the duration of EEF development but allowing enough cell-cell variation in sensitivity to cues to ensure egress is not too synchronous.

Observations that nuclear division within EEFs is arrhythmic^26,27^, combined with our finding that liver egress is plastic and aligned with host feeding-fasting rhythms in normal hosts, indicate that rhythmicity is established between cytokinesis in EEFs and the first IDC. That this plastic scheduling of egress fails in TTFL-deficient hosts^20^ (Figure 1d) is unlikely to matter in natural conditions where clocks are omnipresent, but understanding how parasite rhythms interact with host rhythms is instrumental to elucidate the underlying mechanisms. Recognising that the liver stage can establish the rhythmic development of *Plasmodium* parasites in the blood has several implications. It provides a framework for interpreting how disturbed daily rhythms in humans and mosquito vectors might modulate malaria transmission. For example, plasticity in the time of day that parasites egress into the blood would considerably mitigate any negative impacts that the ongoing shift in *Anopheles* blood feeding times^39–42^ may have on parasites, compared to blood feeding at night^64^. Elucidating the signalling pathways underlying plasticity in EEF maturation could lead to interventions that curb infections at an early stage. Because the transition from asynchronous karyokinesis to synchronous and timed cytokinesis/egress occurs in EEFs and IDC schizonts^26–30,33–38^, agents that disrupt parasite time-keeping have the potential to impair multiple *Plasmodium* life stages simultaneously.

## Methods

We performed three experiments: the ‘Alignment of egress’ and ‘Time of egress’ experiments tested whether parasites egress in line with host rhythms, and the ‘Simulated egress’ experiment quantified the fitness consequences of synchronous versus temporally dispersed egress from the liver. Treatment groups are described in the results section, with detailed experimental designs and procedure timelines in Supplementary Figure S1.

### Parasites, hosts and vectors

We used *P. chabaudi chabaudi* genotype AJ for liver and blood stage infections in the **‘**Alignment of egress’ and ‘Time of egress’ experiments and for the blood stage in the ‘Simulated egress’ experiment. We used *P. berghei* ANKA for liver stage infections in the ‘Simulated egress’ experiment because, due to its preference for young RBCs ^65^, *P. berghei* is unlikely to interfere with the replication of *P. chabaudi* IDC stages, and *P. chabaudi* parasites can be separately quantified using quantitative PCR (qPCR,^45,66^). To independently manipulate the hosts’ feeding-fasting rhythms and the rhythms generated by their canonical circadian clock (TTFL-driven rhythms) we used either wildtype (WT) C57BL/6, or *Per1/2* null (KO) mice that are arrhythmic when housed in constant conditions (constant darkness, *ad libitum* feeding)^43^. All mice were mixed sexes, 8–10 weeks old, housed at 21°C, and fed standard RM3 pelleted diet (801700, SDS, UK) with unrestricted access to drinking water supplemented with 0.05% para-aminobenzoic acid^67^. Mosquito vectors were *Anopheles stephensi*, reared at 26°C, 70% relative humidity, in a 12hr light: 12hr dark cycle, with *ad libitum* access to 8% fructose post emergence. Procedures conformed to the UK Animals Scientific Procedures Act 1986 (SI 2012/3039) and were approved by the ethical review at the University of Edinburgh.

### Transmission of parasites to and from mosquitoes

Donor mice received 10^5^ RBCs parasitised with *P. chabaudi* AJ or *P. berghei* intraperitoneally^68^. *P. berghei* donor mice were treated with 125mg/kg phenylhydrazine 3 days prior to *P. berghei* inoculation to stimulate anaemia and subsequent gametocytogenesis. Female *Anopheles stephensi* mosquitoes (3-5 days post emergence) fed on groups of donors, confirmed positive for gametocytes by microscopic examination of Giemsa-stained thin blood smears, on day 14 (*P. chabaudi*) or day 5 (*P. berghei*) post inoculation. Sample sizes for the ‘Alignment’ and ‘Time of egress’ (*P. chabaudi*) and the ‘Simulated egress’ (*P. berghei*) experiments were 16, 4 and 6 gametocyte donors and 1700, 300 and 350 exposed mosquitoes, respectively. Exposed mosquitoes were housed at 26°C (*P. chabaudi*) or 21°C (*P. berghei*). We quantified the prevalence of infected mosquitoes from oocyst counts obtained from 0.05% mercurochrome-stained midguts, dissected on day 8 (*P. chabaudi*, ‘Alignment of egress’ n=206/1700, ‘Time of egress’ n=20/300) or day 13-14 (*P. berghei*, ‘Simulated egress’ n=50/350) post infection by mosquito bite (dPI). At 13 (*P. chabaudi*) or 20 (*P. berghei*) dPI, we randomly allocated mosquitoes to 62, 13 or 13 pots, containing 24, 20 or 8 mosquitoes each, of which on average 13.0, 6.2 or 6.2 mosquitoes were infected with *Plasmodium (*based on oocyst prevalence) for the ‘Alignment of egress’, ‘Time of egress’ and ‘Simulated egress’ experiments, respectively.

Liver infections were generated by exposing anaesthetised (500µg/kg medetomidine, 50mg/kg ketamine hydrochloride, intraperitoneally) hosts to one pot of mosquitoes each for 20 minutes, after which anaesthesia was reversed using 2.5 mg/kg of atipamezole subcutaneously^68^. Residual *P. chabaudi* sporozoite loads in the salivary glands, at the day of transmission, were quantified by qPCR^8,69^ to estimate sporozoite exposure. Blood stage infections in the simulated egress experiment were established by intraperitoneal injection of 3*10^5^ *P. chabaudi* AJ ring stage-infected RBCs at 39 HPI and additional injections of PBS at 31 and 47 HPI, or by three doses of 10^5^ each at 31, 39 and 47 HPI^68^. To ensure the presence of ring stage-infected RBC at each time point, *P. chabaudi* AJ infected donor hosts were kept at the same photoschedule as the WT hosts, or at photoschedules that were advanced or delayed by 8hrs.

### Perturbation of rhythms and sample collection

Feeding-fasting rhythms were perturbed by restricting food (RF) availability to 12 hours to shift the feeding window in WT mice or create a feeding-fasting rhythm in the KO mice in DD ^12,44^. All mice were acclimated to their photoschedules and feeding-fasting regimes for at least two weeks prior to *Plasmodium* infections. Restricted feeding does not result in weight loss^18^, which we confirmed in the ‘Alignment of egress’ experiment by comparing weights between *ad libitum* (AL) fed and RF groups before infection (F_1,29_=2.00, p=0.168). During all experiments, blood was collected from the tail vein to quantify RBC densities by Coulter counter (2µL, Beckman Coulter) and parasites (10µL for qPCR^70^).

### Molecular protocols

DNA was extracted from 10µL blood using a semi-automatic Kingfisher Flex Magnetic Particle Processor and MagMAX^TM^-96 DNA MultiSample Kit (Thermo Fisher Scientific), following^70^, except for doubling the sample volume. *P. chabaudi* IDC stages were quantified by qPCR targeting the 5’ region of the PSOP1 gene (PCHAS_0620900)^66^, and an AJ-specific section of the 3’ region of the same gene in the ‘Simulated egress’ experiment^45^. PCR efficiency did not differ across plates for the ‘Alignment of egress’ (93±2%, F_4.51_=0.16, p=0.96), ‘Time of egress’ (95±2%, F_3,36_=1.42, p=0.25) or ‘Simulated egress’ (99±2%, F_3,48_=1.66, p=0.19) experiments. No amplification was detected in negative (water) controls. Quantification was relative to a serial dilution of microscopy-counted *P. chabaudi* AJ IDC stages, with a correlation of r^2^>0.99 between parasite densities and PCR quantification cycles for all experiments. We set the lower detection limit to the DNA lowest dilution of the standards, at 5 par/µL.

Residual sporozoites in the salivary glands were quantified as described in^8^. In brief, blood-fed mosquitoes were bisected^71^ and the head-thorax portions were pooled in groups of 3-6 from the within the same pots to extract DNA ^8^. Non-exposed mosquitoes were processed as negative controls. Sporozoites were quantified by qPCR targeting the 18S rRNA gene^69^, using a serial dilution series of *P. chabaudi* DNA from IDC stages, assuming one genome per sporozoite. Amplification efficiency was 93±2.7%, with a correlation between parasite genomes and PCR quantification cycles (r^2^) of 0.99. We approximated sporozoite exposure by summing sporozoite counts for all pools of fed mosquitoes for each individual mouse.

### Statistical Analysis

Data analysis was performed using R version 4.5.0 (2025-04-11). We used linear models to compare PCR efficiencies across plates, and to compare mouse weights and sporozoite exposure (square root transformed) between treatment groups. Infection prevalence was compared between treatment groups using generalised linear models with a binomial distribution. To compare how log_10_ parasite densities changed over time (HPI or dPI, treated as a factor to allow for non-linearity) between treatment groups, we constructed generalised linear mixed effects models, with mouse identity as a random effect to account for repeated measures (“glmmTMB” R package). Models used a Gaussian error distribution, with added zero inflation in the alignment of egress experiment where earlier time points were more likely to be negative. Only mice confirmed to be infected during at least one time point were included. Model assumptions were tested using “Dharma” package simulations. Model selection was performed via model fit (AICc), or log-likelihood ratio test (LRT) for nested models. Tests statistics and associated p-values are presented in the main text. All figures represent the mean ± sem of data.

## Acknowledgements

We thank the BVS staff for animal maintenance; and Alíz Owolabi, Mary Westwood, Jacob Holland, Catherine Oke and Ronnie Mooney for practical help. This work was supported by the Wellcome Trust (grant no. 202769/Z/16/Z; 227904/Z/23/Z), the Royal Society (grant no. URF\R\180020), and the University of Edinburgh (studentship to AHM).

## Author contributions

Conceptualization, SER and AHM; Investigation, AHM, AOD and PS; Analysis, AHM, PS; Writing-Original Draft, AHM, PS and SER; Writing-Review and Editing, all authors; Supervision, PS and SER.

## Data availability statement

The datasets supporting the conclusions of this article are available in the Edinburgh DataShare repository: URL inserted upon acceptance

---

**Figure S1.**
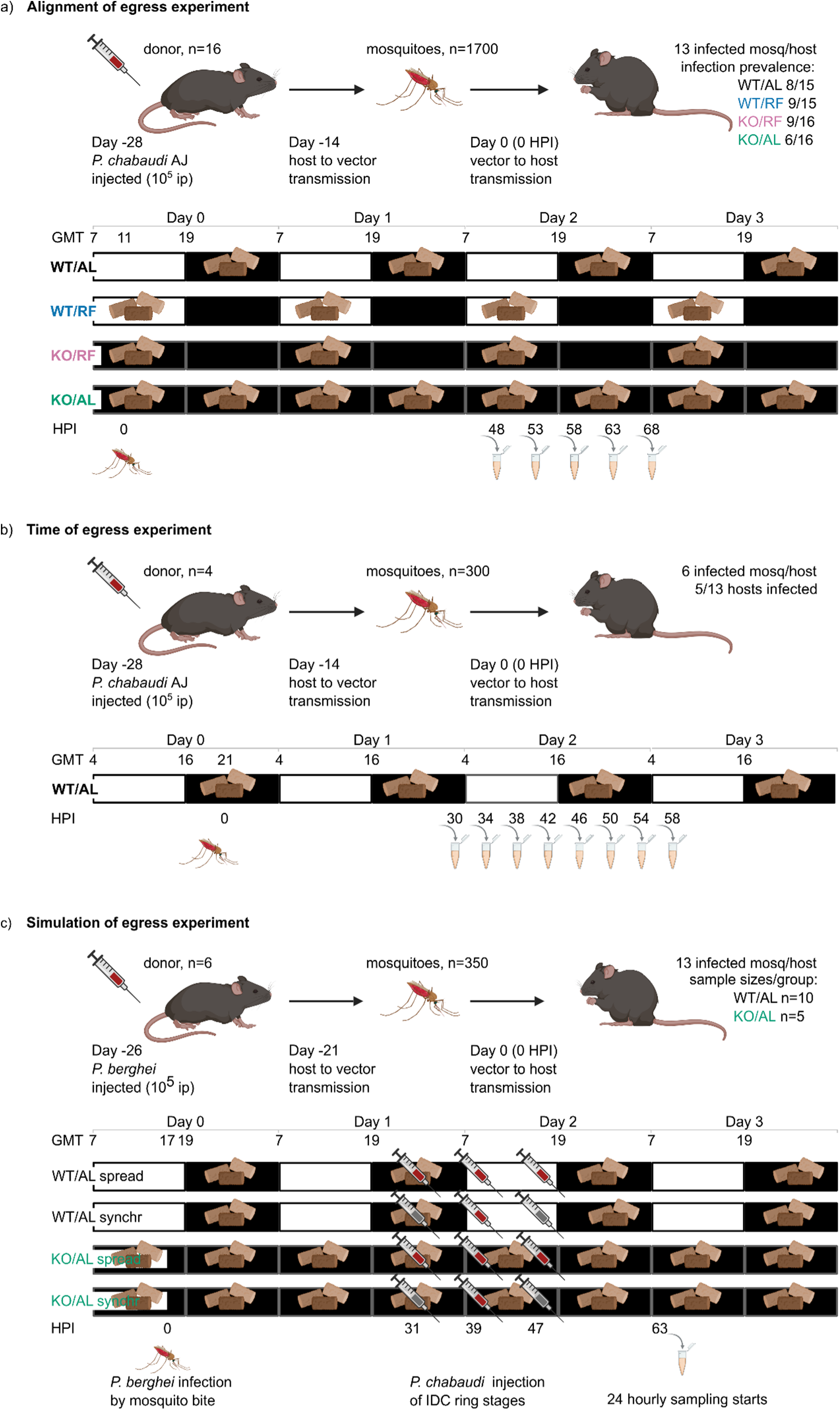
Experimental designs. **a)** Alignment of egress experiment; **b)** Time of egress experiment; **c)** Simulated egress experiment. The top section of each graph depicts the timeline for infection of donor mice, infection of mosquitoes and subsequent infection of mice by mosquito bite, with experimental groups and sample sizes shown. The lower section of each graph illustrates the photoschedules (white: light; black: darkness) and feeding-fasting schedules (pellets: feeding window) for all treatment groups. Sampling time points are shown by illustration of tubes, and injection needles depict the intraperitoneal (ip) injection of *Plasmodium* parasites (red) or PBS (grey). WT: wildtype mice; KO: *Per1/2* null mice; AL: *Ad libitum* fed hosts, eat in darkness for WT and continuously for KO mice in continuous darkness; RF: Restricted feeding for 12 hours, creates a feeding-fasting rhythm in the *Per1/2* null mice in continuous darkness^12,44^; HPI: hours post infection by mosquito bite; GMT: Greenwich Mean Time.

